# DNA Methylation-Based Cell Type Deconvolution Reveals the Distinct Cell Composition in Brain Tumor Microenvironment

**DOI:** 10.1101/2025.01.19.633794

**Authors:** Feili Liu, Jin Qian, Chenkai Ma

**Affiliations:** Department of Neurosurgery, Huashan Hospital, Shanghai Medical College, Fudan University, Shanghai 200040, China; National Center for Neurological Disorders, Huashan Hospital, Shanghai Medical College, Fudan University, Shanghai 200040, China; Shanghai Key Laboratory of Brain Function and Restoration and Neural Regeneration, Shanghai 200040, China; Neurosurgical Institute of Fudan University, Shanghai 200040, China; Shanghai Clinical Medical Center of Neurosurgery, Shanghai 200040, China; Division of Digestive and Liver Diseases, Department of Medicine and Herbert Irving Comprehensive Cancer Research Center, Columbia University Medical Center, New York, NY, 10032, USA; Integrated Diagnostics, Human Health, Health and Biosecurity, CSIRO, Westmead 2145 NSW, Australia

**Keywords:** DNA methylation, tissue deconvolution, brain tumor, cell composition, tumor microenvironment

## Abstract

**Background:** Central Nervous System (CNS) tumors have sophisticated tumor microenvironment (TME) with different cell types such as astrocytes, microglia, neurons, vascular endothelial cells and immune cells. These non-cancerous cells orchestrate the brain TME to regulate cancer progression and therapeutic response. This study aimed to develop a cell composition deconvolution method for CNS tumor and to determine the impact of these cell compositions on patients’ outcomes.

**Methods:** We identified the cell type-specific CpG loci using the pairwise differential methylation analysis for 13 major cell types in CNS. Using non-negative least squares (NNLS) methods, we established this cell-type deconvolution approach, MDBrainT, for brain tumors. The predictive accuracy of our MDBrainT model was tested in the DNA methylation profiling of the purified cell samples and compared against another algorithm. Cell composition was predicted by MDBrainT in several brain tumor (glioma, ependymoma, medulloblastoma and ATRT) cohorts and the correlation between cell composition and tumor molecular subtypes and patient outcomes was also assessed.

**Results:** Cell type-specific CpG loci for CNS TME was used to build MDBrainT model. Based on these DNA methylation markers, MDBrainT predicted the cell composition in the TME including tumor cells with high accuracy. Endothelial cells were predominately presented in glioblastomas while the percentage of CD8 T cells wassignificantly higher in ATRT. A substantial difference of cell composition was two molecular groups of posterior fossa ependymoma (PFA vs PFB). A higher percentage of cells in TME was usually associated with worse outcomes.

**Conclusions:** MDBrainT is a robust algorithm for cell composition prediction for brain TME. Cell composition in brain TME is distinct across different pathological types and molecular subtypes.

**Key Points:**

- MDBrainT is a robust DNA methylation-based deconvolution approach for brain tumor microenvironment (TME).
- Different molecular subtypes of brain tumors have distinct cell composition patterns.
- Cell composition in brain TME informs patients of outcomes.

**Importance of this study:** DNA methylation is a cell type specific marker that has been utilized for tumor molecular diagnosis, disease progression and therapeutic monitoring. A DNA methylation-based classifier for brain tumors precisely predicts the molecular subtypes but not the cell composition of tumor microenvironment. Brain tumors are a complex cell mixture where tumor microenvironment is critical for tumorigenesis and therapeutic resistance. Here, we developed a novel deconvolution approach (MDBrainT) to predict cell composition for brain tumors. Our model has revealed the heterogeneity of cell composition between tumor types. In addition, tumors with different molecular subtypes have distinct cell composition. Cell percentage in TME also informs patients of outcomes. The tumor microenvironment including cell composition of each patient may direct the different regimen of precision medicine.

## Introduction

Central nervous system (CNS) tumors are the most lethal cancer type globally[1, 2]. Primary treatment includes the surgical removals of the tumor with minimizing the neurological injury and maintaining the neurological functions post-surgical operations[3]. Emerging immunotherapy including immune checkpoint blockage (ICB) and CAR-T has exhibited its utility in anti-tumor efficacy and prolonging the patients survivals in solid tumor such as melanoma, lung cancer and colorectal cancer[4, 5]. A few clinical trials are now sought to determine the efficacy of immunotherapy including personalized cancer vaccination and ICB for brain tumors, however, these clinical trials fail to meet the expectations[6-8]. Immune cells such as T cells and NK cells have lost anti-tumor functions as demonstrated in *in vitro* studies due to the complexity of tumor microenvironment (TME)[9, 10]. Unlike other solid tumors, brain tumors have unique TME and infiltrated cellular pattern where it has less cancer associated fibroblasts but more neurons and supporting cells in the CNS. These surrounding neuronal cells promote tumor growth by synapses formed with glioma and mediate the immunomodulation[11, 12]. Similarly, microglia and monocyte derived macrophage facilitates immune evasion in glioma via TLR2 activation[13]. Therefore, understanding the cellular biology and component of CNS TME contributes to the success in immunotherapy for brain tumors.

DNA methylation, mainly on the fifth cytosine in the mammalian cells, is a reliable epigenetic marker for cellular identity that indicates the cell of origin[14, 15]. Using the cell type specific methylome, it traces the original sites of metastatic cancer of unknown primary, classifies the heterogeneous brain tumors on molecular subtype levels and predicts the cell composition from admixture such as cell-free DNA[15-18]. A few attempts have been made to establish DNA methylation-based algorithms mainly for immune cell composition and tumor purity prediction[19-21]. By predicting the fibroblast and 7 immune cells composition in tumors, MethylCIBERSORT, a CIBERSORT-based deconvolution method, stratify tumor patients into “immune hot” or “immune cold” subtypes of which have distinct outcomes and survival benefits from immunotherapy[20]. Based on 419 CpG signatures for 11 leukocyte cell types, MethylResolver deconvolutes the TME by providing tumor-infiltrating immune cells and tumor purity[21]. As CNS has unique microenvironment and these algorithms fail to include neuronal and other supporting cells information in their cell signature references, the utility of these existing algorithms for CNS tumors would likely have less accuracy. Recently, Zhang et al developed a deconvolution approach including several major brain cell types for cell proportion prediction and demonstrated the applicability in aging and brain-related disorders but not in CNS tumors[22].

Here, we developed a DNA methylation deconvolution method for pan-brain tumors (MDBrainT). Based on the cell type specific markers, MDBrainT gives the cell proportion of tumor cells, immune cells, neuronal and supporting cells for bulk brain tumor samples. By benchmarking the discovery cohort, MDBrainT has demonstrated a high predictive accuracy and outperformed other DNA methylation deconvolution methods. We also showed that the utility of MDBrainT for understanding the cellular heterogeneity of TME across different pathological types and molecular subtypes of brain tumors, which is beneficial for informing patients therapeutic management.

## Methods

### Cell type and brain tumor specimen datasets

To establish the major cell type specific DNA methylation signature in brain, we include these following cell types: astrocyte (n = 7), oligodendrocyte (n = 24), neuron (n = 43), microglia (n = 17), dura (n = 10), vascular endothelium (n = 4). In addition to the major brain cells, we also recruited immune cells including B cell (n = 6), CD4+ T cell (n = 6), CD8+ T cell (n = 6), NK cell (n = 6), neutrophil (n = 6) and granulocyte (n = 6) to recapitulate the tumor microenvironment in brain. Detailed information regarding the source of the cell type reference datasets is listed in **Supplementary table 1**.

We subset dataset covering 10 common cancer types (Astrocytoma, ATRT, DMG, Ependymoma, Glioblastoma, Low Grade Glioma, Medulloblastoma, Meningioma, Oligodendroglioma, Subependymoma) in brain from a publicly available cohort of brain tumor DNA methylation profiling [15]. This dataset was split into two sub-datasets according to the author labels (Training and Validation cohorts). To evaluate the performance of MDBrainT and assess the utility of cell composition for brain tumor patients’ outcomes, a few additional datasets were included.

### Signature selection

For cell type signature selection, we first identified the differential DNA methylation CpG loci in a pairwise approach after consolidating all 147 purified cell DNA methylation profiles by removing the CpG loci located in the sexual chromosomes or cross-reaction CpG loci. CpG loci with adjusted p value < 0.01 and |log Fold Change|> 2 was regarded as differential CpG loci. After the duplicated loci were removed, the remaining 26095 CpG loci was taken into the next step. Then, we performed a differential methylation CpG loci analysis (|log Fold Change|> 2) for all tumor samples as compared to the non-tumor controls. Lastly, we removed the duplicated CpG loci between cell type signatures and tumor signatures and some CpG loci with missing values. A total of 25913 CpG loci was considered as brain tumor microenvironment cell type signatures (**Supplementary Table 2**). Cell type reference matrix was generated by aggregating these cell type signatures of DNA methylation profiles with same cell type by the median.

### Deconvolution

To predict the cell composition in any given samples, we performed a non-negative least squares (nnls) analysis, which has demonstrated the performance and utility in a few previous studies. In brief, we first transformed the DNA methylation beta value of each given sample into a matrix with CpG loci in row. Subsequently, we applied “nnls” algorithm on the sample matrix against the cell type signature reference matrix. Predicted cell composition was calculated by the output for each cell type divided by the sum. The cell type predictive accuracy is assessed by RMSE and median of the predicted proportion for the samples having same cell identity. The same approach was utilized for purified cell samples and tumor samples from cohorts. For benchmarking EpiDISH and MethylResolver deconvolution, we used the same cell type signature matrix as the reference. EpiDISH and MethylResolver were performed with default settings.

### In silico simulation

For each cell type excluding tumor samples, we mixed in the DNA methylation beta value profile of every sample from cell type cohort with samples of other cell types in ratios of 0 to 100, 10 to 90, 20 to 80, etc. up to 100 to 0. Afterwards, we applied MDBrainT to the admixture to predict cell composition of this in silico admixture. The consistency of predicted versus actual cell composition at each admixture ratio was visualized in the boxplot with the diagonal reference line.

### Tumor purity

Tumor purity was determined by the tumor cell composition yield by MDBrainT. For benchmark purposes, we also calculated the purity predicted by the “InfiniumPurify”. Correlation coefficient of predictions by these two algorithms were assessed by the Pearson correlation test.

### Survival

Proportional hazard model was adapted to test whether the cell composition was associated with the brain tumor patients’ overall survival (OS) and progression-free survival (PFS) using Log-rank test. High or low cell percentage categories of each cell type were determined by the optimal cut-off estimation.

### Statistical significance

The statistical significance of two groups was examined by Wilcoxon test. Pearson correlation analysis was adapted for the correlation significance. P value less than 0.05 was considered as the statistical significance.

## Results

### Development of MDBrainT

To develop a DNA methylation-based cell type deconvolution algorithm for CNS tumors, we first identified the major cell type specific markers from the published datasets. We collected the DNA methylation profiles from 147 purified cell samples, which covered the major cell types (astrocyte, oligodendrocyte, neuron, microglia, dura, endothelial cell, B cell, CD4 T cell, CD8 T cell, NK cell, Neutrophil and granulocyte) in the brain. Through pair-wise comparison, we selected the statistically differently expressed CpG loci for all normal cell type specific markers. For the DNA methylation-based markers in tumor cells, we selected the hyper- and hypo-methylated CpG loci as compared to the normal brain tissues with variety of status. After the overlapped loci of normal cell types and tumor specific being removed, a total of 25913 loci were regarded as the cell type specific markers for brain tumors (**Figure 1B, supplementary table 3, supplementary figure 1**). Of these markers, most were located in the Open Sea (62.03%) and gene body (68.09%) (**Supplementary figure 2**).

**Figure 1.**
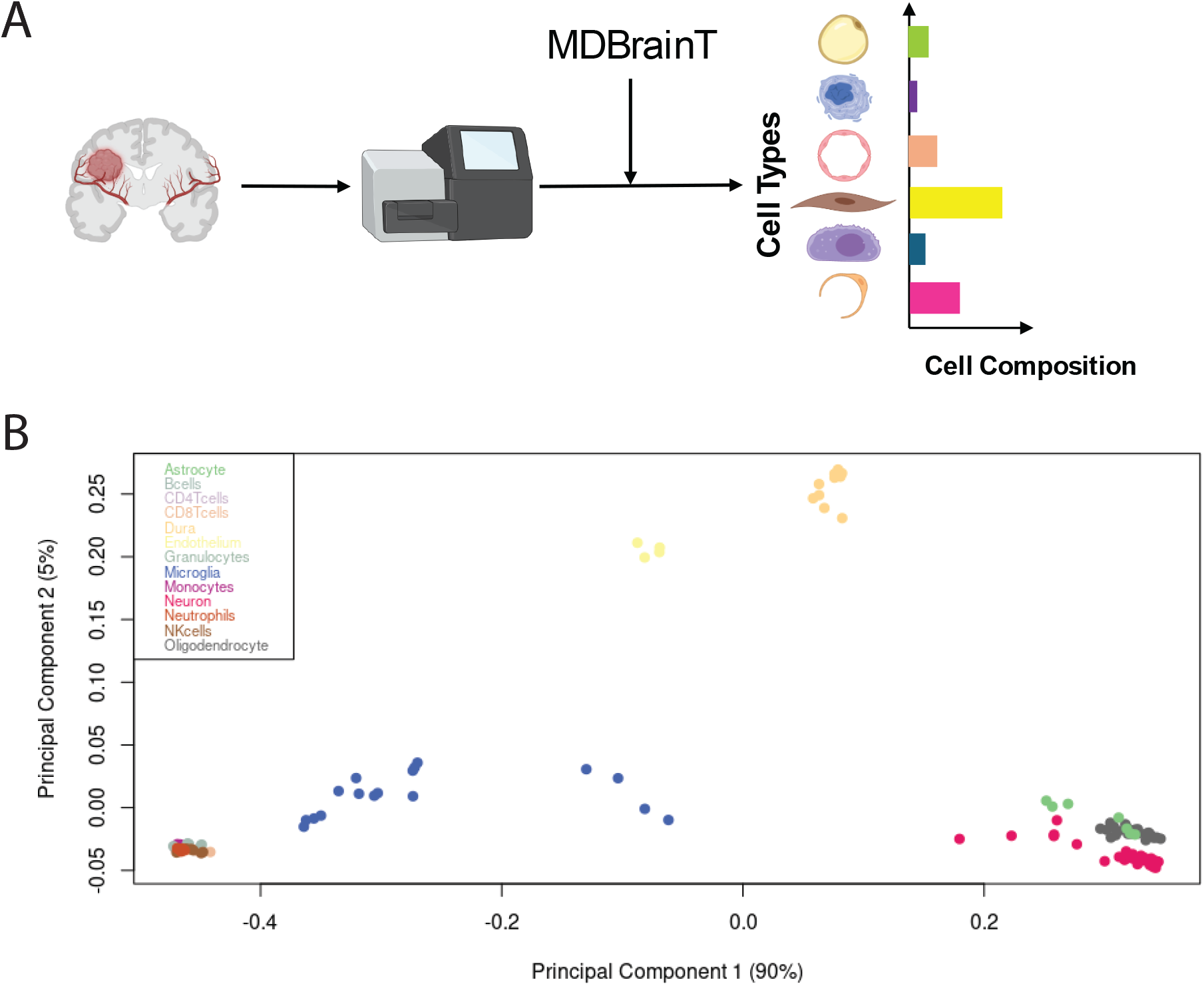
Development of MDBrainT using DNA methylation profiling of major cell types in brain. **A**, Schematic workflow of cell type deconvolution for brain tumor using DNA methylation profiling. **B**, PCA plot showing the global DNA methylation profiling of the discovery cohort.

Next, we sought to test the predictive performance using our markers. We first tested our brain tumor microenvironment cell type prediction model in the discovery cohort. The predicted cell type was consistent with the actual cell types in the majority of samples (139/147) (**Figure 2A**). RMSE also suggest the average prediction errors of our model were relatively low (< 0.1)(**Figure 2B**). One CD4 T cell sample was classified as CD8 T cell and one astrocyte sample was misclassified as oligodendrocyte. Only four astrocyte samples (4/12) and two oligodendrocytes (2/25) were assigned to the neuron classification. That is likely because of the low purity of the sample sources or the proximity of these cell types. Astrocytes with the highest RMSE were likely to be misclassified into other cell types. Then, an *in-silico* simulation was performed to test the accuracy of percentage prediction of each cell type. A computational mixture of known compositions of the target cell type mixed with all other cell type for the rest composition were adapted to compare the predictive composition with the actual composition. As we expected, our model yielded an accurate prediction especially when the compositions of cell mixture were below 80% (**Figure 2C**). To comprehensively evaluate the performance of MDBrainT, we subsequently directly compared the predictive accuracy of MDBrainT against EpiDISH and MethylResolver using our cell type signatures. More off-target cell types were identified using EpiDISH and MethylResolver as compared with MDBrainT. As expected, our MDBrainT (median accuracy: 0.91995) outperformed EpiDISH (median accuracy: 0.85739) and MethylResolver (median accuracy: 0.89284) in terms of the average of cell composition prediction for 13 major cell types (**Supplementary figure 3**).

**Figure 2.**
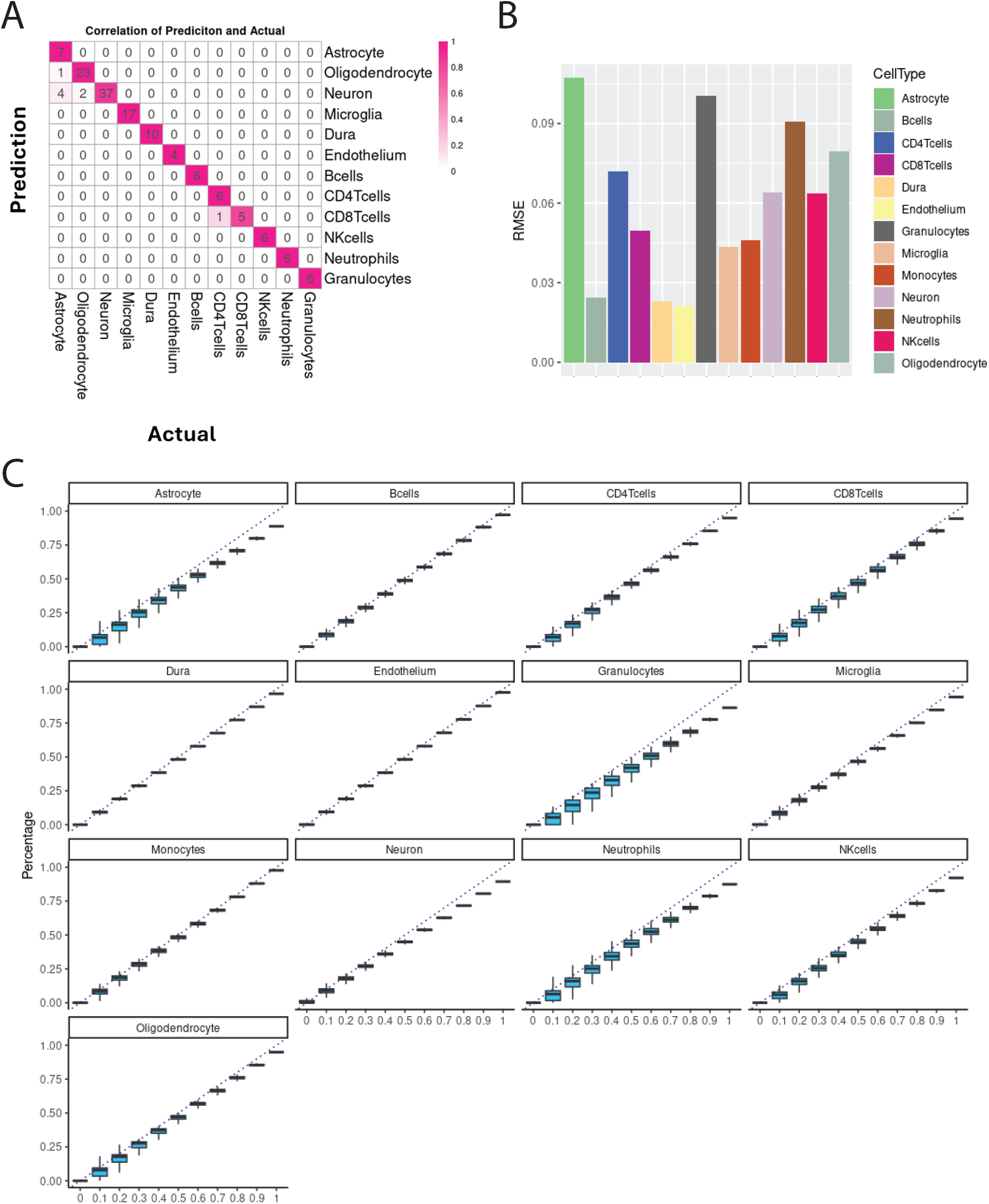
The performance of MDBrainT algorithm. **A**, Confusion matrix showing the predicted cell types versus the actual cell type in the discovery cohort. **B**, Barplots showing the RMSE for each cell type. Black dot line indicates the average of RMSE. **C**, in silico simulation showing the consistence of predicted cell composition versus the actual cell composition from 0% to 100%.

### Different Brain Tumors have distinct cell composition patterns

To determine the variance of cell type composition in the different types of brain tumors, we then applied our model in a large cohort that profiling the DNA methylation of 1756 cases of most common brain tumors, including Astrocytoma, AT/RT, Diffuse Midline Glioma (DMG), Ependymoma, Glioblastoma (GBM), Low-Grade Glioma, Medulloblastoma, Meningioma and Oligodendroglioma. A large variance of cell composition was identified across different brain tumors. High grade brain tumors (GBM and Medulloblastoma) usually contain higher tumor purity, as compared with low grade tumors (LGG and Meningioma) (**Figure 3A**). These results were consistent in the validation cohort **(Figure 3B**). We also benchmarked the consistency of our model’s tumor purity prediction with previous model. Although our model indicated less tumor purity prediction in the low-grade brain tumors, there was a significant correlation between our model and InfiniumPurity, except for ependymoma (**Figure 3C, Supplementary Figure 4**). All brain tumor types had a significant correlation between MDBrainT and MethylResolver regarding tumor purity. In both correlation analyses, DMG and GBM had a relatively low correlation suggesting the intertumoral heterogeneity. MDBrainT also had substantially less time to run as compared to EpiDISH or MethylResolver (Supplementary Figure 4). Notably, microglia presented in the low-grade brain tumors such as LGG and meningioma while vascular endothelial cells were enriched in the high-grade brain tumors like GBM and ATRT. For GBM, methylated MGMT patients had a remarkably higher vascular endothelial cells composition (**Figure 3E**). There was no statistical significance of the endothelial cell composition between the primary and recurrent GBM (**Figure 3F**). High level of vascular endothelial cells was also recapitulated in another three GBM cohorts, suggesting the robustness of our MDBrainT. Female GBM patients had a significantly higher proportion of vascular endothelial cells than male GBM patients (**Supplementary Figure 5**).

**Figure 3.**
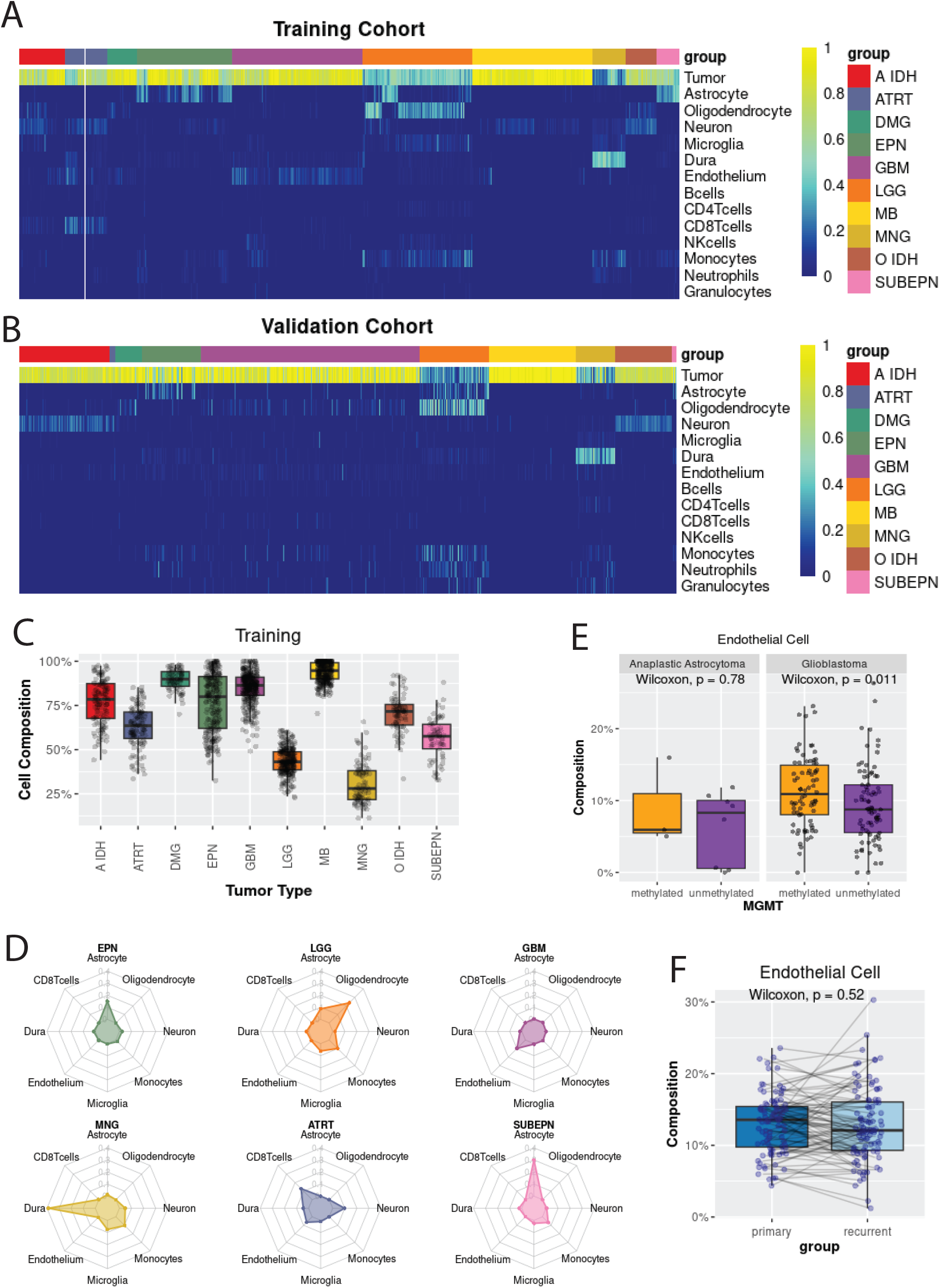
Brain tumors are featured by the distinct cellular microenvironment. Heatmap showing the different cell type composition presents in each major brain tumor types for Capper et al Training cohort **(A)** and Validation cohort **(B). C**, Brain Tumor purity is correlated with the grade of brain tumors. **D**, unique cell composition is recapitulated in the six types of brain tumors. The average predicted cell percentage in each type of brain tumor is plotted in the radar plot.

### Cell compositions reflect molecular subtypes of brain tumor

In addition to the heterogeneity of cell composition across different brain tumor types, large variance of cell composition was also observed within the same brain tumor type. We examined the predicted cell composition in ependymoma, which had the most variance of cell composition across all types of brain tumors. As expected, distinct predicted cell contribution was described in the molecular subtypes of ependymoma. In posterior fossa ependymoma, PFA had lower cell compositions of astrocyte (p = 0.00034), microglia (p = 5.9*10^-9), vascular endothelial cells (p = 5.2*10^-8), monocytes (p = 1.1*10^-8) but a higher neuron percentage (p = 4.4 *10^-5) as compared with PFB. For supratentorial ependymoma (RELA and YAP), a significantly high cell proportion of microglia (p = 0.032) and monocytes (p = 0.028) was observed in RELA. No difference in cell composition was identified between two spinal ependymoma (MPE and SPINE) except astrocytes (p = 0.00022), monocytes (p = 0.001) and CD8 T cells (p = 0.0022) (**Figure 4A**). To investigate whether the cell composition was affected by age or gender, we then applied MDBrainT model to a PFA ependymoma cohort. No differences in cell composition were found between patients’ gender but a significantly weak correlation of age with cell composition of astrocyte and neuron and tumor purity was identified (R = 0.33, p < 0.001). (**Supplementary figure 6**)

**Figure 4.**
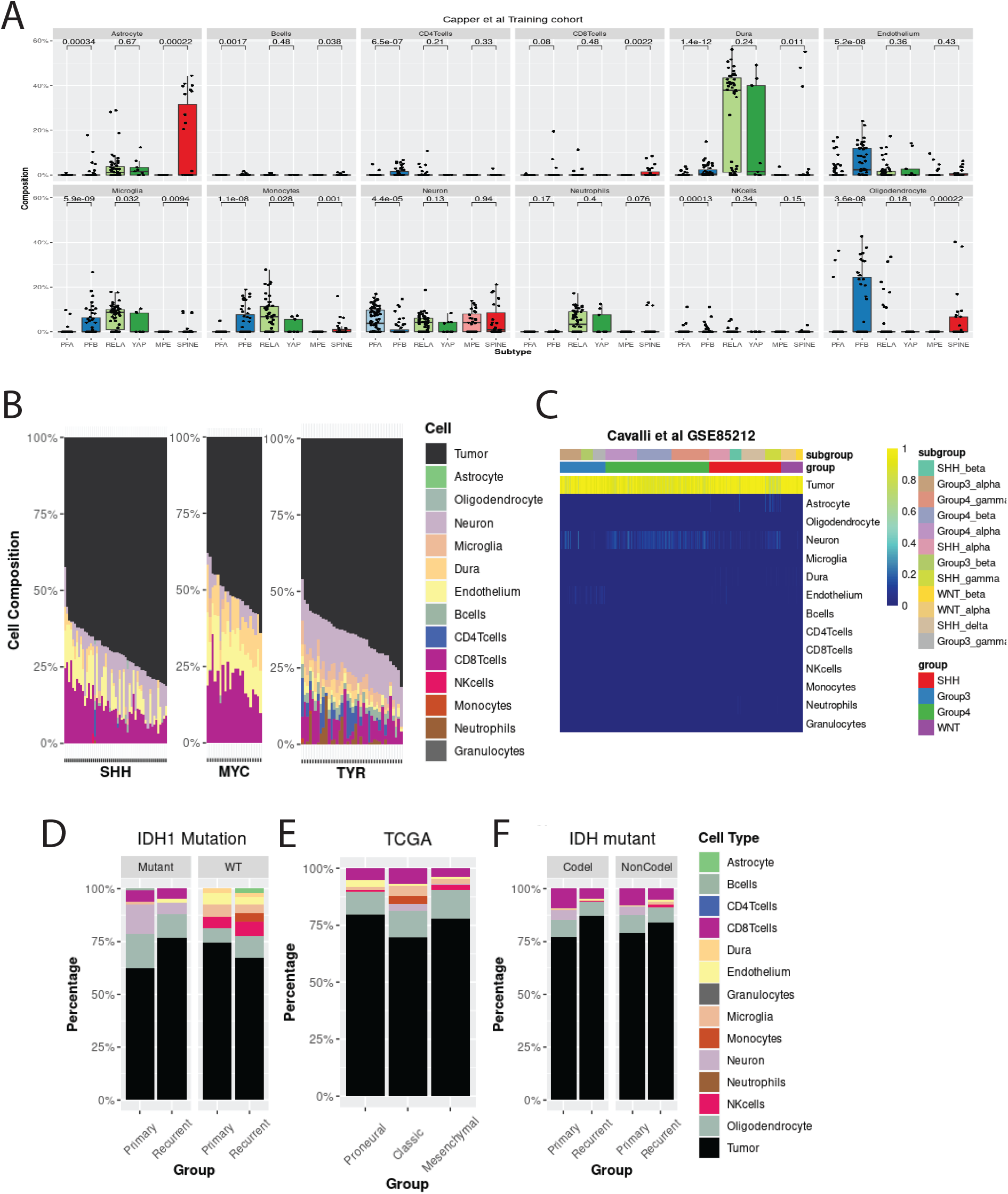
Heterogeneity of cell composition is reflected in the Molecular Subtype of Ependymoma. **A**, Significant difference of Astrocyte, microglia and neuron percentage in the different molecular subtypes of ependymoma with the same anatomical location. **B**, distinct cell composition between SHH, MYC and TYR subtypes in ATRT. Each column represents individual patients. **C**, high tumor purity and less non-cancerous cells was identified across MB subtype and subgroups.

For ATRT, more than 10% of non-cancerous cells were considered as the CD8+ T cells, suggesting the high immune cells infiltration (CD8+ cells) (**Figure 3D and 4B**). Although CD8+ T cells, neuron and endothelial cells were relatively higher in ATRT as compared to other brain tumors, we also observed the heterogeneity of cell composition regarding three molecular subtypes of ATRT. TYR subtype ATRT had predominantly neurons in the microenvironment while MYC subtype ATRT had relatively low percentage of neurons. Instead, CD8+ T cells and endothelial cells were the majority of cell types in the tumor microenvironment of SHH and MYC subtype ATRT. (**Figure 4B**) For MB, WNT MB had the least healthy cell composition of all four molecular subtypes. Consistent with previous results (**Figure 3A**), MB had a relatively high level of tumor purity. Neuron and astrocytes were the most common non-cancerous cell types in Group 3 MB. Group 4 MB had a relatively higher level of neuron than the other three subtypes. SHH MB, especially SHH gamma subgroup, had more astrocytes and neurons in the tumor microenvironment. (**Figure 4C**). These results highlighted the distinct cellular microenvironment in brain tumor and personalized cell therapy strategies could be applied to the patients diagnosed as the different molecular subtypes. For glioma, a distinct cell composition was also observed between IDH1 mutant and wild type. In IDH1 wild type glioma, more non-cancerous cell types and a proportion of vascular endothelial cells were identified, likely because more GBMs were categorized as IDH1 wild type glioma. In contrast, a higher percentage of neurons were found in IDH1 mutant glioma, especially in primary tumors (**Figure 4D, Supplementary figure 7**). We also explored the cell composition across three transcriptomic subtypes in GBM. Classica GMB exhibited a higher percentage of non-cancerous cells (**Figure 4E**). Stratifying IDH mutant gliomas by 1p/19q co-deletion (Codel) revealed primary or recurrent IDH1 mutant gliomas had similar cell composition regardless of 1p/19q co-deletion status (**Figure 4F**). Taken together, these results indicated distinct cellular TME would be taken into consideration for immunotherapy or cell therapy when patients had different molecular subtypes.

### Cell composition informs the patient outcomes

To further validate that the cell composition indicated of the patients outcomes with ependymoma, we then investigated the patients OS and PFS stratified by the cell percentage. In PFA ependymoma, higher cell composition of astrocyte in TME was associated with the worse OS and PFS (Log-rank test, p = 0.0099 and 0.01, respectively). Although higher proportion of astrocytes was associated with the worse survival in RELA ependymoma, statistical significance was identified in PFS (Log-rank test, p = 0.00066) not in OS (Log-rank test, p = 0.34)(**Figure 5A**). Similarly, a higher composition of the endothelial cells in GBM is associated with the OS (Log-rank test, p = 0.049) but not the PFS (Log-rank test, p = 0.13) (**Figure 5B**). These results highlighted the cellular TME orchestrated the malignant progression in CNS tumor.

**Figure 5.**
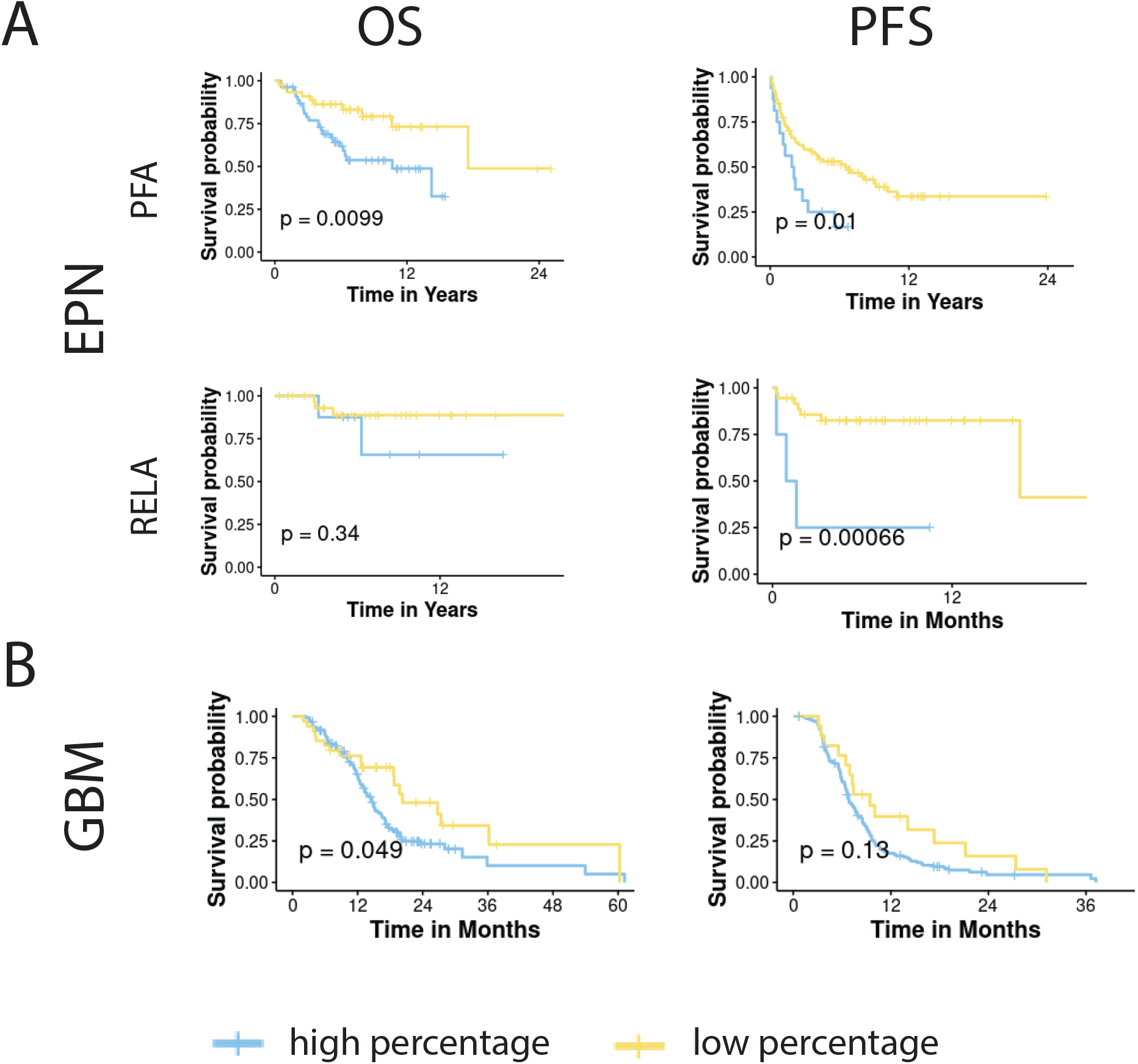
Cell composition is associated with patient OS and PFS. **A**, OS and PFS of PFA and RELA EPN, stratified by the astrocyte proportion. **B**, OS and PFS of GBM, stratified by the endothelial cells proportion.

## Discussion

Cellular composition in brain tumor TME has drawn the attention recently because it proves these surrounding cells like neurons, astrocytes and microglial cells have critical role in suppressing the function of immune cells and promoting the tumor growth[23]. Our DNA methylation-based deconvolution for cell composition in brain tumors provides an insight into the brain TME. Distinct cell composition in the brain TME suggests that different cell therapy or immunotherapy strategies would be considered after the surgical operation or the first-line treatment failures.

Precise biomarker for cell types and a suitable mathematic model are the keystone for cell type deconvolution. A pair-wise comparison could reduce the bias of one-versus-others approach where the average values of all other off-target samples are likely masked by most samples. A few transcriptome-based models have demonstrated their utility in the solid tumors for TME and tumor infiltration T cells prediction, which indicates the immunotherapy benefits[24, 25]. DNA methylation-based classification enables precision tumor diagnosis, highlighting the importance of DNA methylation in brain development and high specificity of methylated CpG loci as biomarker for brain tumors[15]. A few attempts have also been made to establish DNA methylation-based deconvolution approaches for liquid biopsy and minimal residual diseases detection[26, 27]. Benchmarking studies have revealed that there is not much difference in predictive accuracy using transcriptomic data though Machine Learning based algorithm, CIBERSORT, is slightly better than others[28]. However, DNA methylation-based signatures are more reliable markers than transcriptomic signatures for glioma because it reflects the cell identities for not only tumor cells but also the TME[29]. Adding DNA Hypersensitive Site information into partial correlation analysis benefits the accuracy of DNA methylation-based deconvolution model and performs better than CIBERSORT[30]. Our results reveal that MDBrainT outperforms EpiDISH and MethylResolver when the identical DNA methylation CpG loci signature is utilized as cell type specific markers, suggesting that deconvolution results are also affected by different mathematical algorithms and NNLS is a robust reference-based interface for cell type prediction using DNA methylation data.

Tumor purity varies between tumor types and samples and can be inferred by copy number alteration, transcriptomic expression and DNA methylation[21, 31, 32]. A benchmarking assessment of 10 copy number- or gene expression-based tumor purity algorithms reveals a large discrepancy of predicted tumor purity results across different approaches[33]. Since DNA methylation is an epigenetic marker reflecting the cell identity during tissue development and DNA methylation-based cell type deconvolution exhibits high accuracy of tumor purity prediction, emerging attempts to decipher solid tumor purity using DNA methylation [20, 34-36]. MDBrainT extends the utility of DNA methylation-based deconvolution into CNS tumor. For non-cancerous cell types, we also include in CNS specific cell types such as astrocytes, neurons and dura, which overcome the limitation of previous methods are immune cells only. InfiniumPurity doesn’t pre-define the non-cancerous markers or limits the non-cancerous markers for cell type deconvolution. Instead, it needs to provide non-cancerous tissue samples as the normal reference, which may not be able to achieve in brain tumor specimens. To overcome this, MDBrainT also includes in normal cell types as reference matrix so no normal tissue was required for performing cell composition prediction. We observe that tumor purity predicted by MDBrainT is slightly lower than that inferred by InfiniumPurity, which is likely because there are non-immune cell types are included in the MDBrainT reference.

TME is one of the determinants for tumor progression and the success of novel immunotherapy and cell therapy[37]. Therefore, investigating TME by cell composition prediction provides a better understanding of the cellular TME and interaction of the non-neoplastic cells and tumor cells. Different to other organs, CNS has unique TME and cellular composition. Our study provides a landscape of cell composition in brain tumor TME. Similar to the results from scRNAseq data, a non-neglectable proportion of non-cancerous cells are identified in brain TME[38, 39]. Myeloid-derived cells such as macrophage coordinate the proliferation and cytotoxicity of CD8 T cells in glioma during tumor progression [39, 40]. To accommodate the glioma tumorigenesis and progression, astrocytes promote the angiogenesis by releasing cytokines and growth factors while BDNF-TrkB signaling in neuron-glioma synapses is activated to amplify the glutamate intake via AMPA receptor on the malignant cells[41, 42]. Patients with high neural content in tumors are also associated with decreased OS and PFS in a large cohort[29]. High level of neurons in TME is observed in astrocytoma, oligodendroglioma, AT/RT and medulloblastoma, suggesting that blockade of neuron-tumor interaction is likely a potential therapeutic strategy.

Numerous studies elucidate the genetic and epigenetic profiles across brain tumor molecular subtypes or TME in one certain type of brain tumor, but few explore the differences of TME between molecular subtypes yet. Using snRNAseq, Hoogstrate et al. describes a reduction in tumor purity and endothelial cell but increases of neuron and oligodendrocyte in the recurrent IDH1 wild type glioma, which is consistent with our results[37]. Based on transcriptomic data, IDH1 wild type GBM can be categorized into three distinct TME subtypes. Longitudinal analysis also demonstrates that the dynamic changes of TME subtypes during GBM recurrence. Notably, this TME subtyping informs of the patients’ response to the immune checkpoint blockage[43]. Although these studies highlight the importance of TME in brain tumor progression and therapeutic response prediction, they fail to integrate non-immune cell into the cell type prediction and compare the TME across different molecular subtypes. Each subtype of ependymoma has its own cell composition in TME. Even for ependymoma sharing the same anatomic location, the cell composition is not identical. This discrepancy of cell composition also contributes to the different survival outcomes. The distinct cell composition across molecular subtypes is described by our study, highlighting the complexity of TME and different cell-cell interaction across molecular subtypes.

We also notice there are some limitations in this study. Firstly, the cell type specific CpG loci in this study is not necessary in the same genomic region, which might be difficult to get the biological insight on these cell type specific biomarkers, especially when most CpG loci are located in the Open Sea and gene body. Differential methylated region analysis will be helpful to overcome it and shortlist the CpG loci locating together. Secondly, due to the unavailability of purified cell DNA methylation profiling such as pericyte, we cannot generate a whole cell atlas for brain TME though MDBrainT covers the major cell types. Lastly, we did not compare the changes of cell composition between pre- and post-therapeutics such as anti-VEGFR for GBM. Longitudinal studies will be beneficial for uncovering the dynamic changes of cell composition before and during target therapy and immunotherapy.

In summary, we introduced a novel DNA methylation-based cell type deconvolution method tailored for CNS tumor with high predictive accuracy. IDH1 mutant glioma (astrocytoma and oligodendroglioma) have a relatively high proportion of neuron in TME. Different molecular subtypes have distinct cell composition in TME. The proportion of non-cancerous cells in TME is associated with patient outcomes.

## Code and data availability

MDBrainT can be accessed and downloaded from github:

https://github.com/mackaay/MDBrainT

No new data is generated from this study.

## Notes

**Conflict of Interest** There is no conflict of interest between authors.

### Competing Interest Statement

The authors have declared no competing interest.

