## Supplementary figures and images for "DNA Methylation-Based Cell Type Deconvolution Reveals the Distinct Cell Composition in Brain Tumor Microenvironment"

### Supplementaryfigure1.pdf

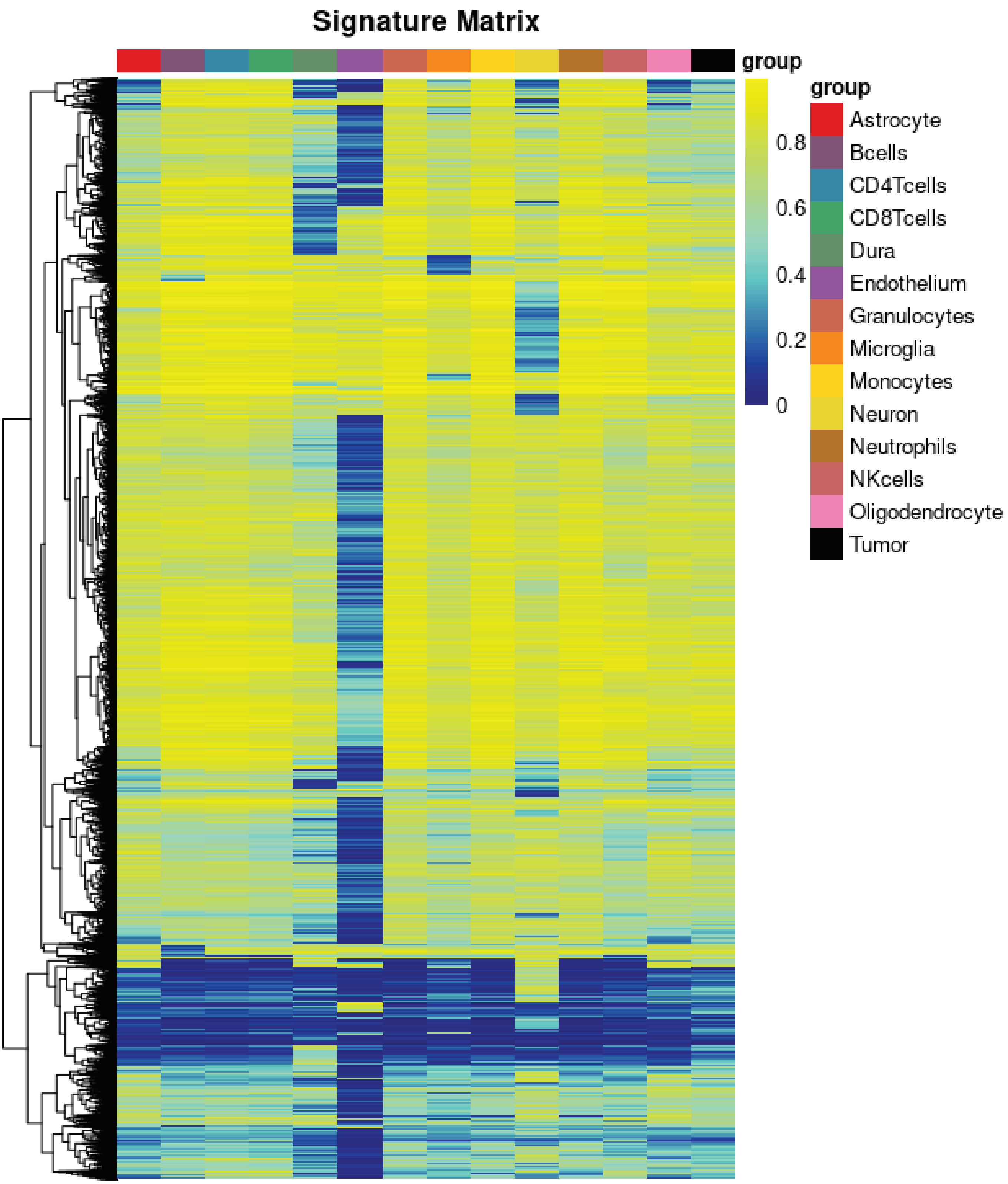

### Supplementaryfigure2.pdf

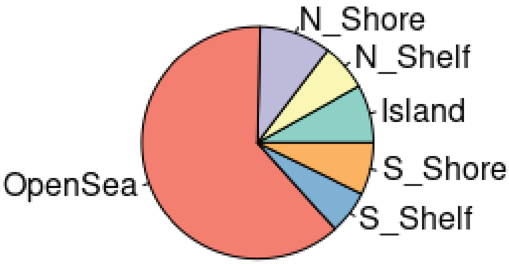

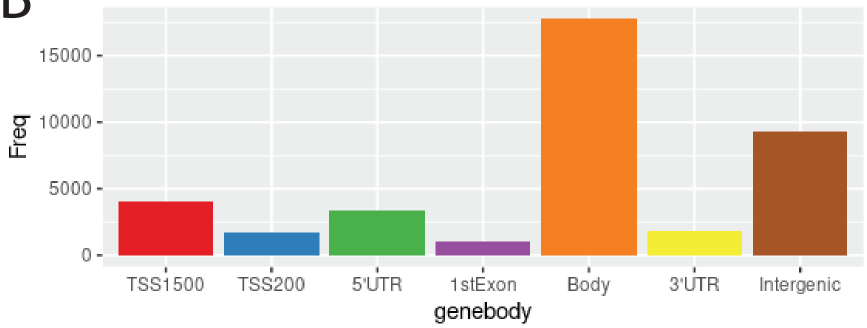

### Supplementaryfigure3.pdf

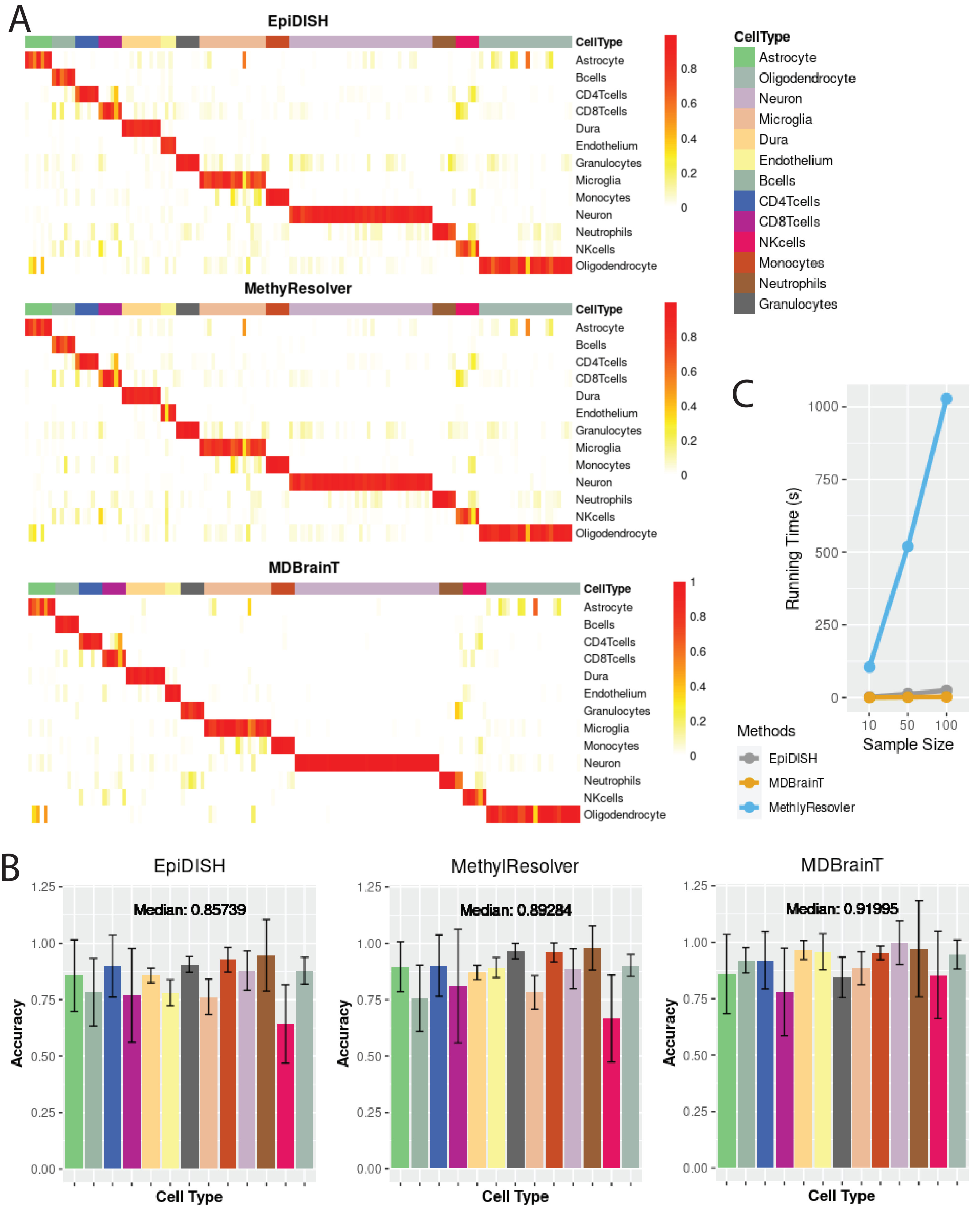

### Supplementaryfigure4.pdf

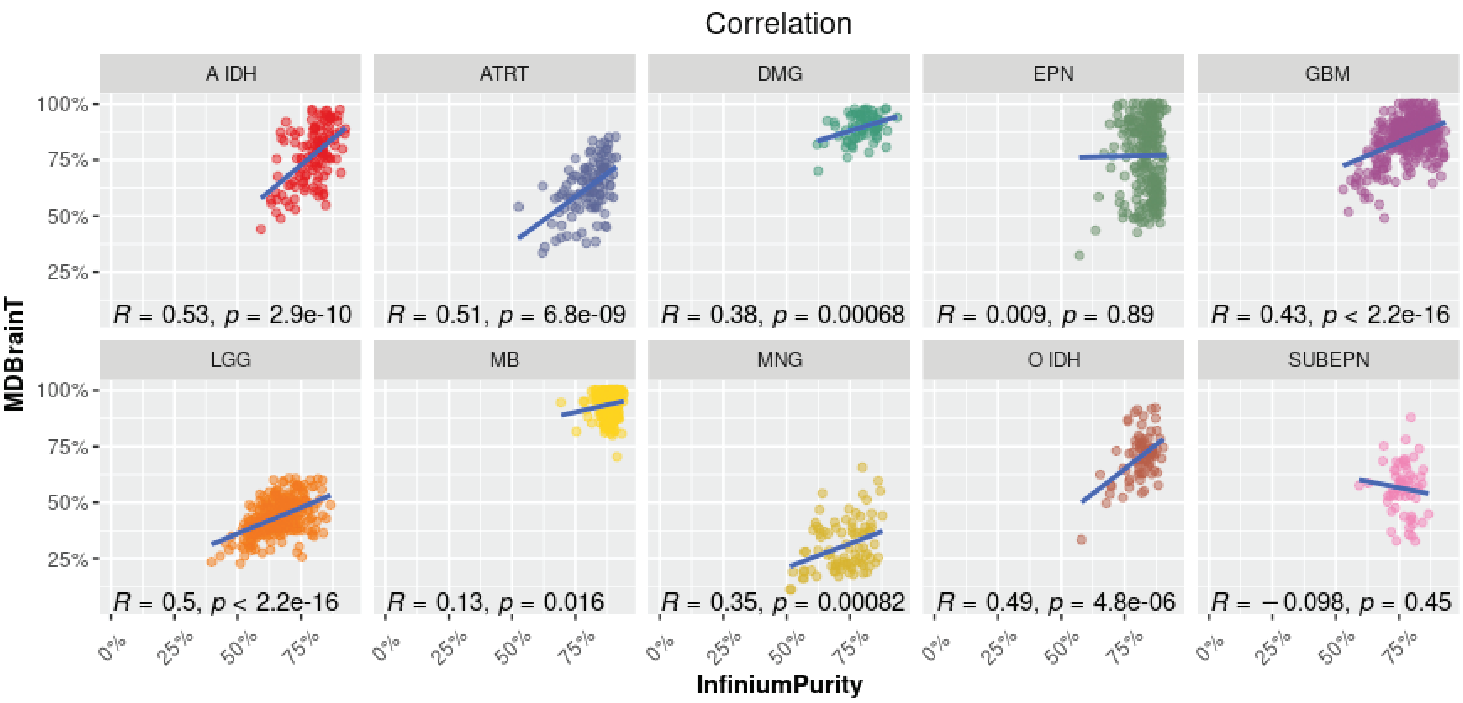

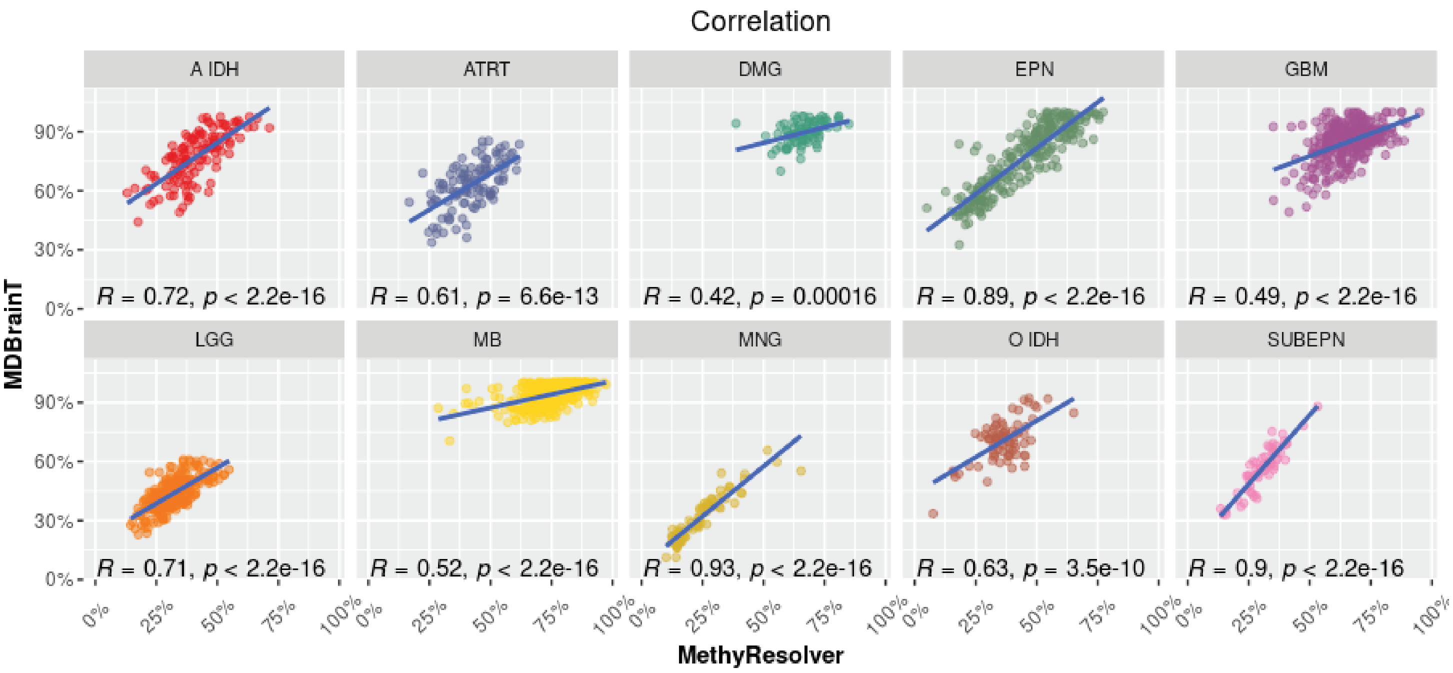

### Supplementaryfigure5.pdf

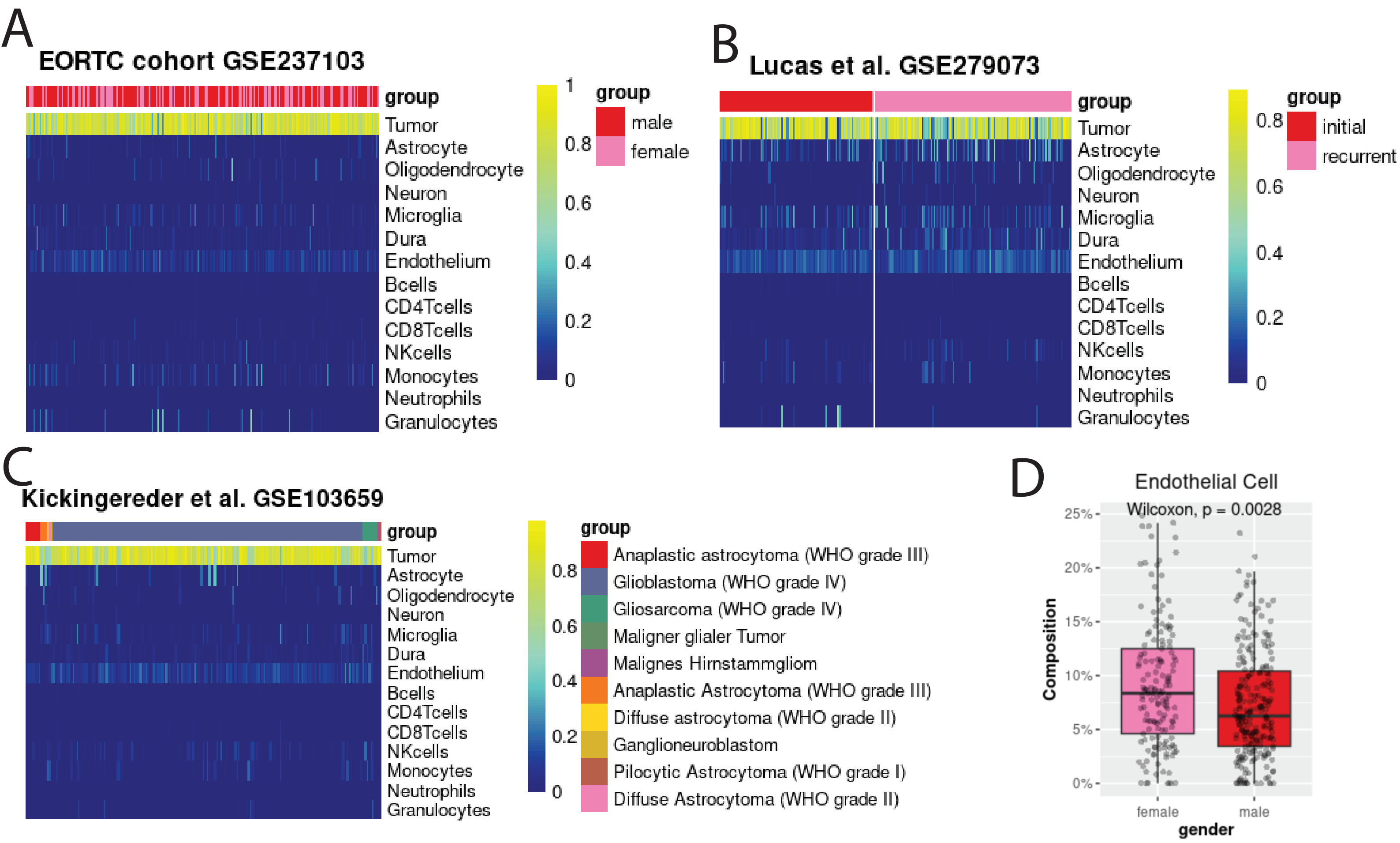

### Supplementaryfigure6.pdf

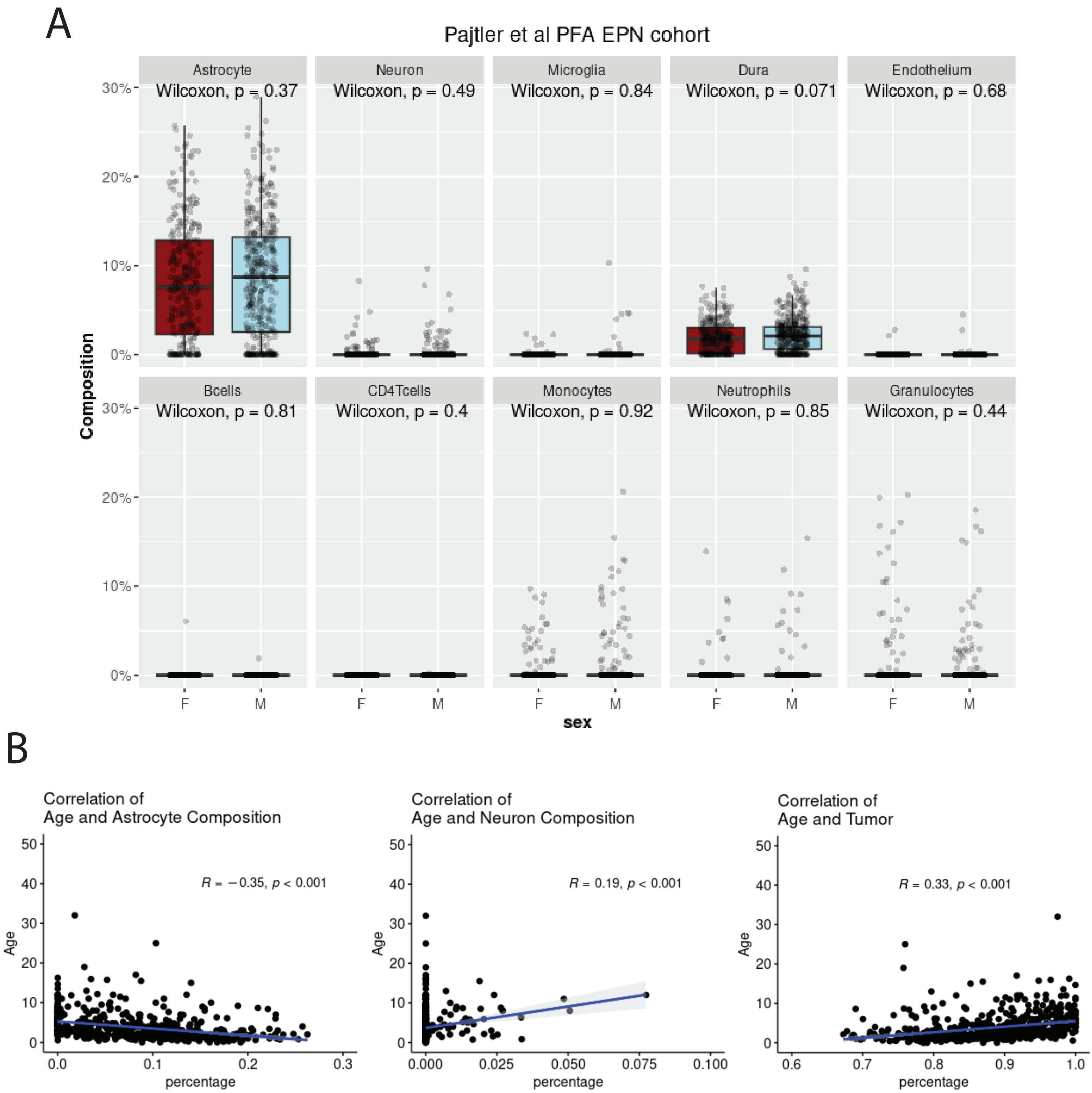

### Supplementaryfigure7.pdf

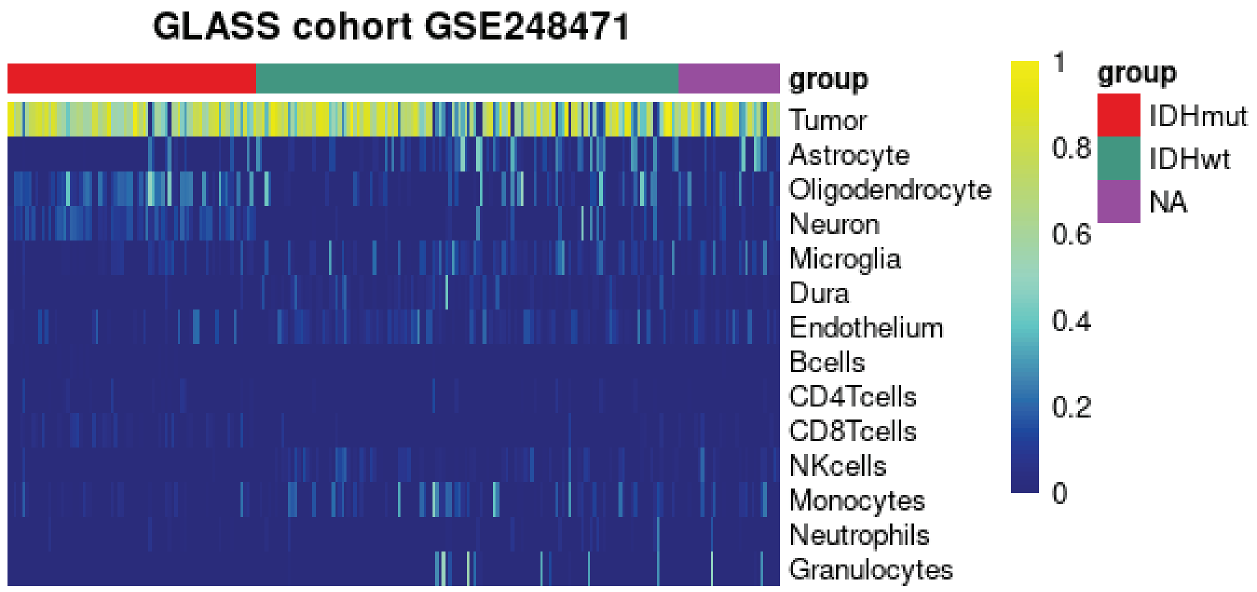
